# Kin selection underpins family dynamics in rural China

**DOI:** 10.1101/2023.09.04.552901

**Authors:** Qiao-Qiao He, Ming-Yang Wang, Jia-Jia Wu, Tian-Jiao Feng, Xiu-Deng Zheng, Jie-Ru Yu, Song-Hua Tang, Ling-Ling Deng, Chang Fu, Ruth Mace, Yi Tao, Ting Ji

## Abstract

The number of co-resident members is an important characteristic of human families and thus plays an important role in human evolution. However, an inclusive fitness model of how many members should co-reside has not been explicitly tested. Here we use an evolutionary game model and a decade of family dynamic data from a rural population of southwestern China (1,110 households) to show that the existence of an evolutionary stable family size can be understood by considering the average local relatedness and average inclusive fitness of the family. Empirical data analyses show that average relatedness 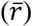 decreased with increasing family size, and 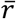 was associated with helping behavior (especially for the elders) in the family, so the family tended to keep a higher level of average relatedness by adjusting the number of co-resident members. These findings are helpful to understand the variation in family size in different cultures and to demonstrate the importance of kin selection in regulating family dynamics.

## 1. Introduction

Human families usually comprise kin groups through which the descent lines are traced; these co-resident adults are responsible for the protection and the physical and psychological nourishment of offspring (Fox, 1983; Gardiner & Bjorklund, 2007). Therefore, kin selection theory is a key to understanding human family dynamics evolutionarily (Hamilton, 1964a, 1964b). As one of the most important structural characteristics, human families vary widely in the number of co-resident members (and in relationships between them) in different historical times and cultures (United Nations Department of Economic and Social Affairs Population Division, 2017, 2019). Thus, studies of the change in family size under the framework of kin selection theory can provide important insight for understanding human evolution.

Large families which used to be common in many regions around the world, are still prevalent in some rural places (Burch, 1967; Stack, 1974) and becoming more important in some developed countries (Bengtson, 2001). Living in a large family is associated with various benefits, such as risk resilience (Dessy et al., 2021a; Sauerborn et al., 1996), more help (Thomas et al., 2018), lower living expenses costs (Chomtohsuwan & Nagashima, 2010a), and workloads (Chen et al., 2023). Although it has been argued that these families seldom exceed 25 persons per household (Burch, 1970). Some existing studies described the determinants such as demographic events (fertility, mortality, life expectancy, age at marriage, etc.) (Burch, 1970; Collver, 1963; Levy, Marion J., 1965), and cultural practices, such as post-marital residences (patrilocal, matrilocal, neolocal, duolocal, etc.) (Brown, 1987; Ember, 1971; Hrnčíř et al., 2020; Porčić, 2010) and marriage systems (monogamy, polygamy, etc.) (Dessy et al., 2021b) that affecting or limiting family size, others emphasized economic limitations (Lang, 1946) or psychological difficulties (Hsu, 1943) of maintaining large family size. However, an inclusive fitness model of how many members should co-reside has not been explicitly tested. Several studies examined the effects of kinship dynamics on behaviour and life history evolution, suggesting that the age-linked change in local relatedness affects menopause and sex differences in helping and harming behaviours (Croft et al., 2021; Ellis et al., 2022; Koster et al., 2019; Mace & Alvergne, 2012), whilst the effects of family size on the change of average local relatedness and consequent cooperation and competition behavior remain unexplored.

Emlen (Emlen, 1995) first applied evolutionary theories to explain the formation, dynamic, and dissolution of human families and the interactions between family members. As Emlen (Emlen, 1995) pointed out, when offspring get more inclusive fitness by staying than leaving natal families to reproduce independently, families form; in contrast, when the latter one is bigger than the former, offspring disperse, and families decrease in size and even break up. Immigration or the birth of a new family member results in increasing of family size, and the emigration or death of a former family member causes decreasing in family size (Cohen, 1969). Thus, the family dynamics, and changes in family size and composition, should influence the average relatedness of co-residing members, which may affect the cooperation and conflict between and within households and ultimately individual’s inclusive fitness (Rodrigues & Gardner, 2013a; White et al., 2019). This raises the question of whether there is an evolutionary stable (or optimal) family size concerning a family’s average inclusive fitness under the framework of kin selection theory.

The concept of optimal group size has been well described by behavioralists who studied the evolution of foraging group size (Smith, 1985). Benefits of group living have also been well documented (including foraging, reproductive and protective functions (Whitehouse & Lubin1, 2005)). Family groups share similar group-living benefits with other social groups, plus some extra inclusive fitness benefits by helping co-resident relatives (Hamilton, 1964b). Although insider-outsider conflict theory predicts that an optimal-sized group could be unstable (Sibly, 1983), family groups show higher stability than other sorts of social groups, and last for a considerably longer time than most of them (White et al., 2019).

A recent study suggested that family size was determined by both resource defense benefits from group-defended resources and collective action benefits from the cooperation of group members (Shen et al., 2017). However, we argue that the genetic relatedness of group members might underpin both kinds of benefits, as it decreases social conflict and thereby increases both types of group benefits at any given family size. Of course, the relatedness of family members itself might change with family size, and living in a large family often means a higher probability of interacting with distant or unrelated family members, therefore its effect on group benefits might also change with size. Here we considered how relatedness varies with family size and influences family-group size using an evolutionary game model, and we tested the model predictions using 10-year longitudinal data from a matrilineal human population, the Mosuo. Since the co-reside family members normally exhibit extensive helping and interactions with each other than with those who moved out (Davis & Daly, 1997), we restrict the definition of the human family to include only individuals co-reside with each other (Teachman et al., 2013), and throughout the paper, we use “family” and “household” interchangeably.

The Mosuo of southwest China is one of the few populations living in a duolocal matrilineal social system (where husband and wife live apart) (He et al., 2016; Walsh, 2004; Wu et al., 2013). Traditionally, Mosuo men and women do not leave their natal household for their whole life, and husbands visit their wives at night (visiting or “walking” marriage, also called “zouhun” in Mandarin or “sese” in the Mosuo language). Typically, the Mosuo live in large matrilineal households which once could contain up to 20-30 members (Walsh, 2004), comprised of one or multiple grandmothers, her/their brothers, sons and daughters, and offspring of their daughters. Nevertheless, recruiting female or male family members through adoption or cohabiting is common when households lack descendants of either sex. The Mosuo prefer large households and emphasize harmony among family members (Shih, 2009).

However, the proportion of duolocal residence declined and neolocal residence (couples and children living together and do not reside with other kin) increased in recent years (Ji et al., 2016), and patrilocal and matrilocal households are also recorded in the Mosuo society. Households have diminished in size (see Results for details). Rodrigues and Gardner (Rodrigues & Gardner, 2013b) showed theoretically that temporal group-size heterogeneity has stronger cooperation effects than spatial heterogeneity. Thus, using Mosuo population as an example, we mainly focus on how family-size dynamics during a decade are associated with relatedness changes and demography in this period, plus how within-household cooperation varied with family size.

Firstly, we established an evolutionary game model between family members to investigate how the change in family size, and the benefit and cost of helping affect the average inclusive fitness of the family. Secondly, we measured how the average relatedness of a household varied as group size changed during a decade using the longitudinal demographic data of a matrilineal Mosuo populations. Thirdly, we investigated how the family size and its dynamics influenced dispersal behaviour, female reproduction, and individual mortality, and lastly, we studied how family size affected cooperation within the household, in terms of investments to family in an economic game, and proportions of members worked in the farm. We did not consider male reproduction, as previously we found few associations between natal household characteristics and fertility of men (He et al., 2016; Ji et al., 2013), probably due to the unusual duolocal residence system where the parents of children typically live in different households with the maternal family investing most in children day to day.

## 2. Method

### 2.1. Theoretical thinking: family size and average inclusive fitness

We first considered a family with size *n* (i.e., this family has *n* members). For simplicity but without loss of generality, we here ignore the gender and age structure of the family members. This simplifying assumption provides a possible starting point for revealing the evolutionary mechanism determining the size of a family.

We assume that the relatedness coefficient between any two individuals is *r*, based on the classic definition of inclusive fitness (Hamilton, 1964a, 1964b), when individual *i* interacts with individual *j*, that is, individual *i* pays a cost *c* such that individual *j* receives a benefit *b* with *rb* > *c* (Hamilton, 1964a, 1964b), the inclusive fitness of individual *i*, denoted by ∅_*i*|*j*_, can be given by ∅_*i*|*j*_ =−*c* + *r*_*ij*_*b* for *i, j* = 1,2 …, *n*, where *r*_*ij*_ (with *r*_*ij*_ = *r*_*ji*_) represents the relatedness of individuals *i* and *j* (Hamilton, 1964a, 1964b). Moreover, we assume that individual *i* (*i* = 1, 2, …, *n*) will interact with all other individuals with equal opportunities.

Thus, based also on the additivity of inclusive fitness (Hamilton, 1964a, 1964b; Maynard Smith, 1982), the total inclusive fitness of individual *i*, denoted by ∅_*i*_, can be expressed as 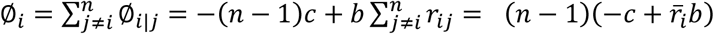 for *i* = 1,2, …, *n*, where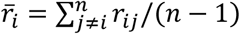 denotes the average relatedness between individual *i* and all other individuals. For convenience, we here use abbreviation TIF to denote the total inclusive fitness of an individual. Therefore, the average TIF of all family members, denoted by 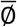, is 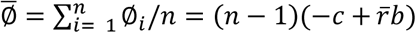, where 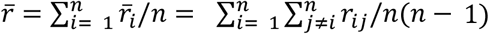 denotes the average relatedness of a family. The average TIF measures the effect of kin selection on the whole household or reflects a possible connection between kin selection and family size.

The average relatedness of a family is determined by family size and the composition of family members. Larger household is more likely to have a more complicated structure, which may have multi-generation or unrelated immigrants, and thus lead to a decrease in the average relatedness of the household. Therefore, without loss of generality, we here define that the average relatedness of the family can be expressed as an exponential function of family size, which is given by 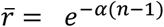, where the parameter α is a positive constant. Based on this definition, the average TIF, 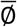, can be rewritten as 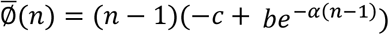.

Note that 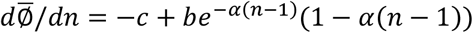 and 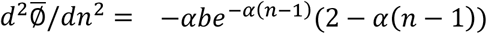. Thus, for α < 1, there must exist a *n*^*^ in the interval 1 < *n*^*^ < (1 + α)/ α such that 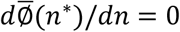 and 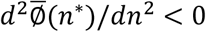, that is, 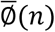 has maximum value at *n* = *n*^*^(see Fig. 1).

**Fig. 1.**
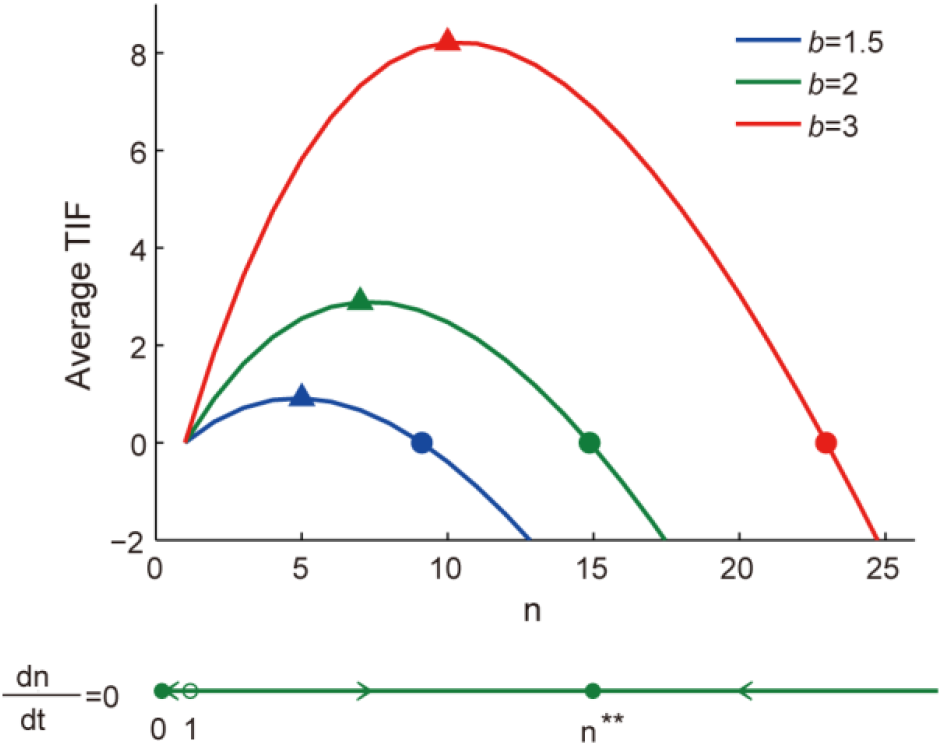
The average TIF (total inclusive fitness) for different household sizes (*n*) and benefit (*b*) between cooperation among relatives, α = 0.05, c = 1. Triangles indicate maximum average TIF, and circles represent stable average TIF. Blue, green, and red lines represent *b* = 1.5, 2, 3, respectively. The line in the bottom shows the interior equilibriums that are stable (0 and n^**^, circle) and unstable (1, hollow circle) when *b* = 2.

### 2.2. Data collection

We used data from three waves of the demographic survey, an economic game involving the allocation of money between oneself and the household, and spot observations on who was working on the farmland. In the years 2007, 2012, and 2017, we conducted demographic censuses every five years for 1,110 households in 17 Mosuo villages in Sichuan Province, China. Most of the inhabitants are Mosuo, and others are Yi, Han, Pumi and a few Tibetan people. As previously described elsewhere (He et al., 2016, 2022; Wu et al., 2013), we visited every household and interviewed an adult representative about both personal information of all family members and household information, and allowed them to refer to other family members on unfamiliar things. We followed changes in 378 Mosuo duolocal households from the year 2007 to 2017, of which most married members applied for duolocal residence in the year 2007. Traditionally, individuals applying for duolocal residence live with their maternal kin, whereas husbands and wives live apart, and children live with their mothers (He et al., 2016, 2022; Walsh, 2004; Wu et al., 2013). Family-group size includes only the current members of a family and does not include individuals who were born in the family but are now permanently living elsewhere. But we counted individuals temporally living elsewhere as students, soldiers, or migrant workers.

In year 2013, we conducted a household game involving money allocation between oneself and the household in each village, as part of a series of economic games. Procedures and some results of other games, such as those of a gift game, have been previously reported elsewhere (He et al., 2016; Mace et al., 2018; Thomas et al., 2018). In a household economic game, we gave each participant 10 yuan (one day’s wage is around 60–70 yuan in this area at that time) and asked them to decide how much money to give to their households (which they knew that would be doubled and delivered to the head of their households afterward), and how much to keep to their own (which would be given to them immediately after the game without being doubled). All analyses were based on a subsample of 137 men and 166 women from 207 duolocal households.

From the year 2011 to 2012, we observed work patterns on some randomly chosen farms during planting and harvesting seasons. We recorded ownership of the lands and the relationship of all the workers to farm owners. All analyses are based on a subsample of farmlands of 244 duolocal households.

The data are not publicly available due to the need to preserve anonymity, but abridged data are available from the corresponding authors upon reasonable request.

### 2.3. Data analysis

#### 2.3.1. Family size and average relatedness

We calculated the relatedness of each pair of individuals with “AGHmatrix” package (Amadeu et al., 2016) in R, based on genealogical data. We also calculated the average relatedness of each individual to other members of the household each year, as well as the average household relatedness at the beginning of each year and after each demographic change (birth, death, and dispersal). There were 412 neolocal, 220 patrilocal, and 100 matrilocal households in 2007, respectively, defined as the residence type applied by most married members at that time. We also analyzed the relationship between average relatedness and family size in these households.

#### 2.3.2. The effect of family size on dispersal and immigration

For analyzing the effects of different family sizes on adult members’ dispersal decisions, we got a longitudinal dataset of 378 households labelled as “duolocal households” in the year 2007. We conducted a discrete-time event history analysis to study the first dispersal of adults out of those duolocal households between the years 2007 and 2017. We used a mixed complementary log-log link model (Singer & Willett, 2003), with household id as a random effect factor, and dispersal (dispersed = 1, stay = 0) as the dependent variable. We measured the first dispersal of every adult who lived in those 378 households from the very beginning, i.e., the year 2007, and individuals joining these households afterwards were excluded in this analysis, but included to calculate household size and average relatedness after their joining. We also excluded individuals born after the year 1989 and person-years after individuals reaching 61 years old, as the former did not reach 18 years old in the year 2007, and elder Mosuo people rarely disperse from their households (only 14 cases happened in 4,066 person-years of individuals aged 61 – 98 years old).

We also tested how family-group size influenced immigration with a simple mixed linear model, where the number of new group members who joined the duolocal households in a giving year was the dependent variable, the size of the family group at the beginning of the year was the fixed factors, and household id was the random factor. We combined family sizes ≤ 3 and ≥ 15 because families with size smaller than 3 or larger than 15 in some years were rare.

#### 2.3.3. The effect of family size on reproduction decisions of women

To analyze how family size affects the reproduction of female members, we measured the annual probability of birth of reproducing-age females in these 378 duolocal households from the year 2007 to 2017. We conducted two mixed complementary log-log link models (Singer & Willett, 2003), where families larger or smaller than the mean size were analyzed separately. In the models, giving birth or not in a specific year was the dependent factor (yes = 1, no = 0), and female id was a random effect factor. Group size, females’ age, and the number of alive offspring were included as time-variant fixed factors. Models included females who were born from the year 1968 to 1999. Mosuo females were allowed to have up to three children (government’s family planning policy in force since the early 1980s), thus person-years with three alive offspring were excluded (n = 6,696 person-years, 713 females; excluded: 1,053 person-years of 109 females). In models, females’ family sizes at the beginning of the year varied from 2 – 19, and we also combined family sizes ≤ 3 and ≥ 15.

We also tested how group size influences birth interval, age of first birth, alive offspring number, and co-residing offspring number of reproducing age females (age = 15-50) of 378 duolocal households in 2007 (all family groups were duolocal families at that time). We modelled the relationship between these variables and family size in a series of linear regressions and Poisson regressions, with the age of females controlled, and sample sizes varied with specific regressions. We also combined family sizes ≤ 3 and ≥ 15. Analyses of birth interval only involved females who ever had more than one child.

#### 2.3.4. Family size and mortality

We conducted two mixed complementary log-log link models (Singer & Willett, 2003) to analyze the relationship between mortality and family-group size, where families larger or smaller than the mean size were analyzed separately. In the models, survival was the dependent variable (dead = 1, alive = 0), and household ID was a random factor. In fixed factors, group size (with size ≤ 3 and ≥ 15 combined) and age (centered by mean) were time-variant variables, and sex was a time-invariant variable.

#### 2.3.5. Family size and within-household cooperation

Finally, we tested the relationship between family size and within-household cooperation level, measured by the proportion of members who worked on the farm, as well as the contribution to the household by members. We conducted two linear models of the proportion of members observed working as owners on farmland and group size, where groups with size ≤7 and ≥ 7 were analyzed separately. We combined group sizes≤ 3 and ≥ 15 in the models. For contribution to the household in the household economic game, we fitted a multilevel model for analysis of the allocation of money between oneself and the household, with the amount of money given to the household as the dependent variable, and villages and households as hierarchically structured random effects. The fixed variables were sex, age (centered by mean), education year (centered by mean), household size (centered by mean), average relatedness to other household members (centered by mean), and some interactions. Continuous variables involved interaction effects were centered by mean. For the graph, we standardized continuous parameter estimates over 2 s.d. and mean-centered binary estimates to enable comparison of continuous and binary variables (Gelman, 2008). Parameter estimates reported in the main text are unstandardized unless stated.

All statistical analyses were conducted in R v. 4.0.5 (R Core Team, 2022). We used “glmer” and “lmer” functions in “lme4” package (Bates et al., 2015) for a mixed model of event history analyses and a multilevel linear model, respectively. Standardization of model estimates was done by “arm” package (Gelman & Su, 2021). Figures were done with the “predict” function and “ggplot2” package (Wickham, 2016). Models and graphics from models use MATLAB.

## 3. Results

### 3.1. The evolutionary stable family size

Based on the above assumptions and definitions in Method 2.1, the time evolution of family size can be expressed approximately as 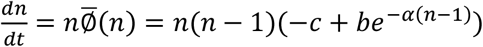. Note that for *n* > 1, the solution of 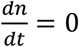, denoted by *n*^**^, can be given by 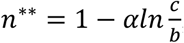, and that *n*^**^ must be stable since 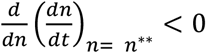. Therefore, *n*^**^ can be called as an evolutionarily stable family size based on the kinship (Fig. 1). Although this simple theoretical model only represents a highly simplified thought experiment, it provides a new perspective for understanding the effect of kin selection on the size of family.

### 3.2. Average relatedness and household size in real human populations

The average family-group size had been slightly decreasing from the year 2007 to 2017. In the year 2007, 378 duolocal households were comprised of 1,520 males and 1,485 females, and the mean size was 7.95 (range 1-18, s.d. = 2.84, mode = 7). In the following ten years, mean size slightly decreased each year (mixed effect model, estimate (ES) = -0.063, 95% confidence intervals (CI) = [-0.073, -0.053], with household ID as a random factor, Fig.s S1-S2), and eventually dropped to 7.33 (range 1-18, s.d. =2.96, mode = 6) in the year 2017. We found no difference in age and sex structure between family groups of different sizes though (Chi-square test, χ^2^ = 52.395 -73.549, and p = 0.112 -0.747 for each year, group size ≤ 3 and ≥ 15 combined, Fig. S3).

The average relatedness of Mosuo duolocal households is 0.345 (s.d. = 0.093, n = 378 households) in the year 2007. As Fig. 2b shows, average relatedness decreases with the size of the duolocal household, and this pattern holds in all 11 years (Fig. S4). Whilst average relatedness increases first and declines later as the size of the other three types of households increases (Fig. 2 and Fig. S4).

**Fig. 2.**
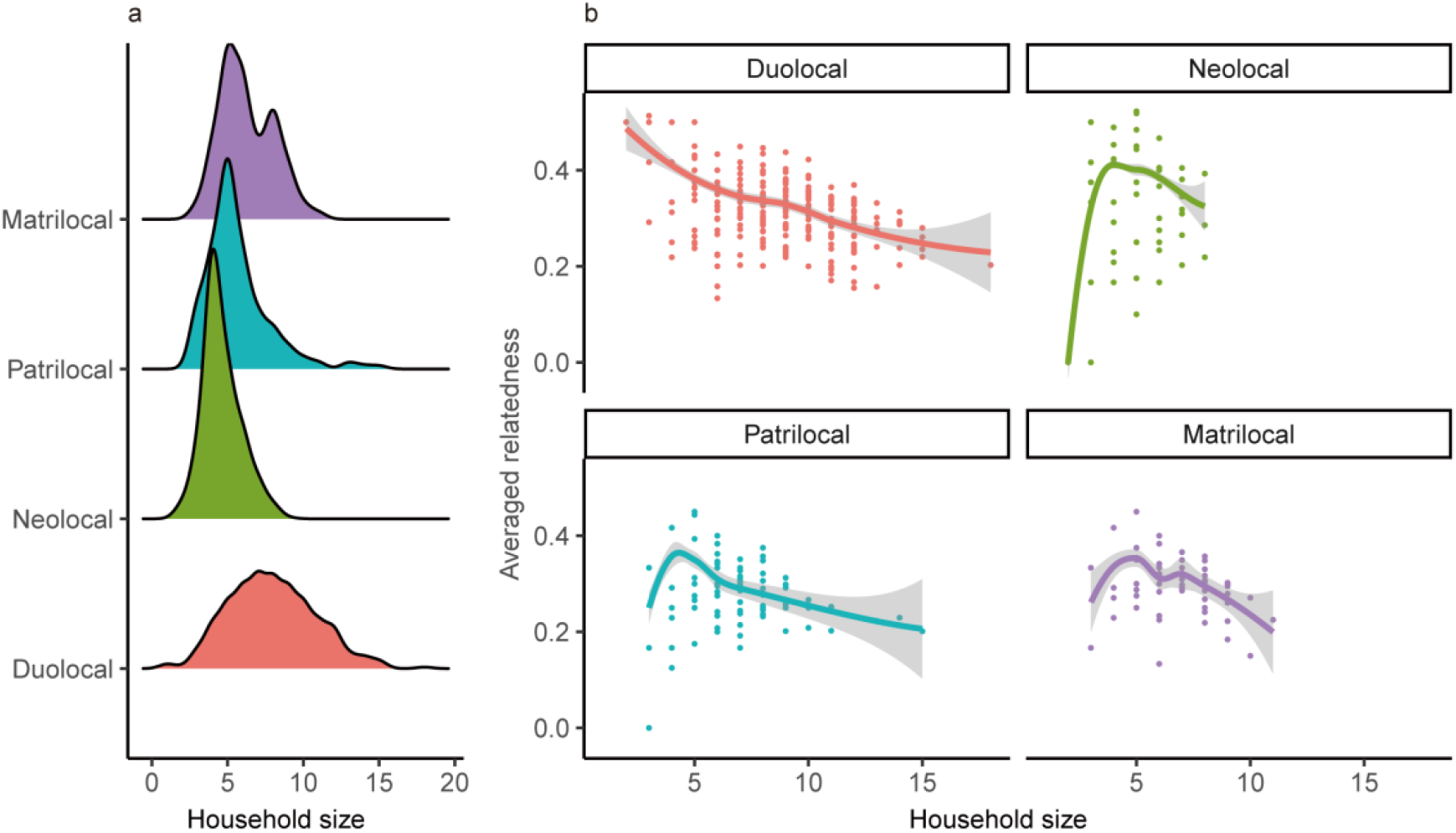
The (a) distribution and (b) the relationship between averaged relatedness and household size in 2007. The average relatedness of families decreased with duolocal household size (n = 378), and it goes up first and then down when household size increases in patrilocal (n = 220), matrilocal (n = 100), and neolocal (n = 412) households. Besides Mosuo, the patrilocal and neolocal data also contain some other ethnic groups (Han, Yi, Pumi, etc.) living in the same villages. Curves fitted using loess. Shaded areas represent 95% confidence interval.

Moreover, we measured how the average relatedness of duolocal household changes after each of four types of demographic events that affected family-group size (Cohen, 1969), of which arrival and birth of a member added one to the size of a household, whilst departure and death of a member minus one of it. In total, 1,111 events happened during the year 2007 and 2017, with the number of emigrations, births, deaths, and immigrations ranked from highest to lowest (Table S1). We found that, on average, household relatedness increased after each departure and death of a member, and in contrast, it decreased after each arrival or birth of a new member (Fig. 3 and Table S1, Kruskal-Wallis test, χ ^2^ = 328.02, p < 0.001). New birth caused less decrease in average household relatedness than joining of new member (post hoc Dwass-Steele-Critchlow-Fligner test, p < 0.001), indicating lower relatedness of new migrants than the newborns to other members of the household. Furthermore, the increase of relatedness caused by death was significantly more than that by leaving (Dwass-Steele-Critchlow-Fligner test, p = 0.008). In total, these results confirm that, the size of households is negatively related to average relatedness.

**Fig. 3.**
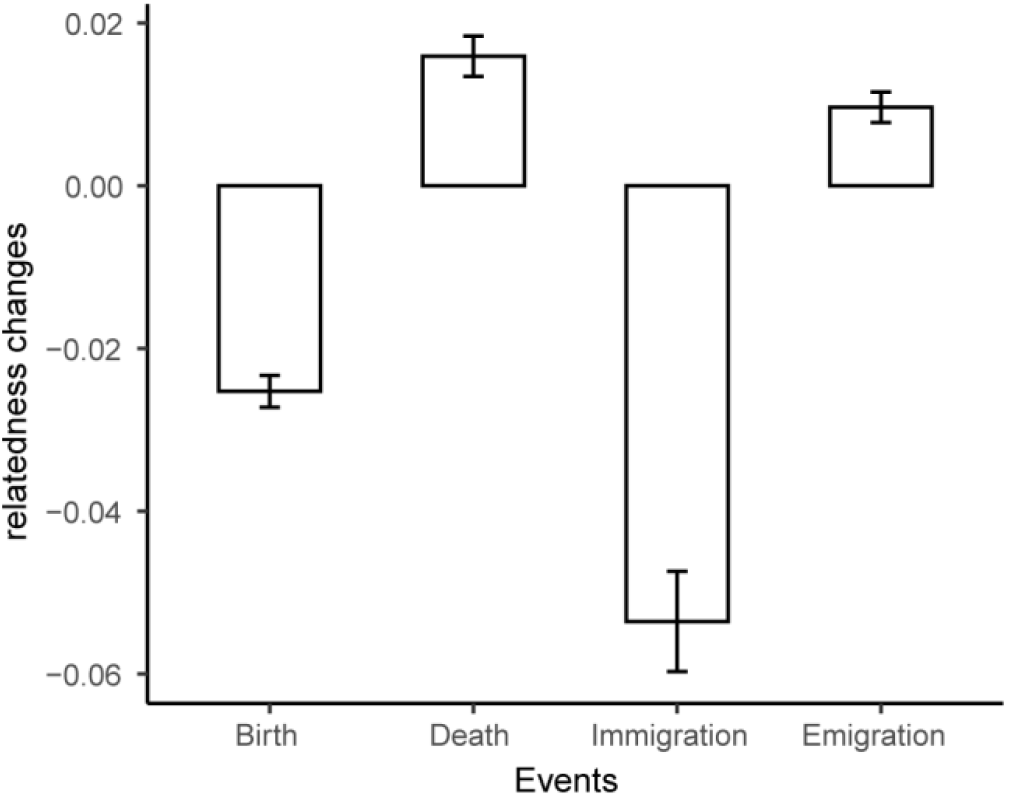
Changes in average household relatedness after each kind of demographic event (one event only involves one person here). Birth and joining of a duolocal household caused the increase in average household relatedness, while death and departure were followed by decreasing in relatedness (Kruskal-Wallis test, χ ^2^ = 328.02, p < 0.001). The Post hoc Dwass-Steele-Critchlow-Fligner test shows that there were significant differences between changes of average relatedness after each pair of comparisons (p = 0.008 for differences between emigration and death, and p < 0.001 for other comparisons). Error bars indicate the standard error from the mean.

### 3.3. Family dynamics in matrilineal Mosuo populations

In total, there were 275 emigrations happened in 15,967 person-years, where 145 women and 130 men (age = 18 -60) dispersed out of their duolocal households (n = 378 households). The theoretical model predicts an arising dispersal probability after household size reached 8. We conducted a mixed discrete-time model of the dispersal decision of adult members from 378 duolocal households between the year 2007 and 2017, and we found some support for this prediction. The hazard of dispersal for adults in families larger than 7 was 1.9 times of those in smaller families (hazard ratio (HR) = 1.884; 95% CI = [1.357, 2.668]; Fig. 4 and Table S2). Moreover, individuals highly related to other members in households showed a lower hazard of dispersal (HR = 0.189; 95% CI = [0.047,0.793], Table S2). We also found that less immigration happened in larger duolocal households (ES of group size = -0.003; 95% CI = [-0.006, -0.001], Table S3).

**Fig. 4.**
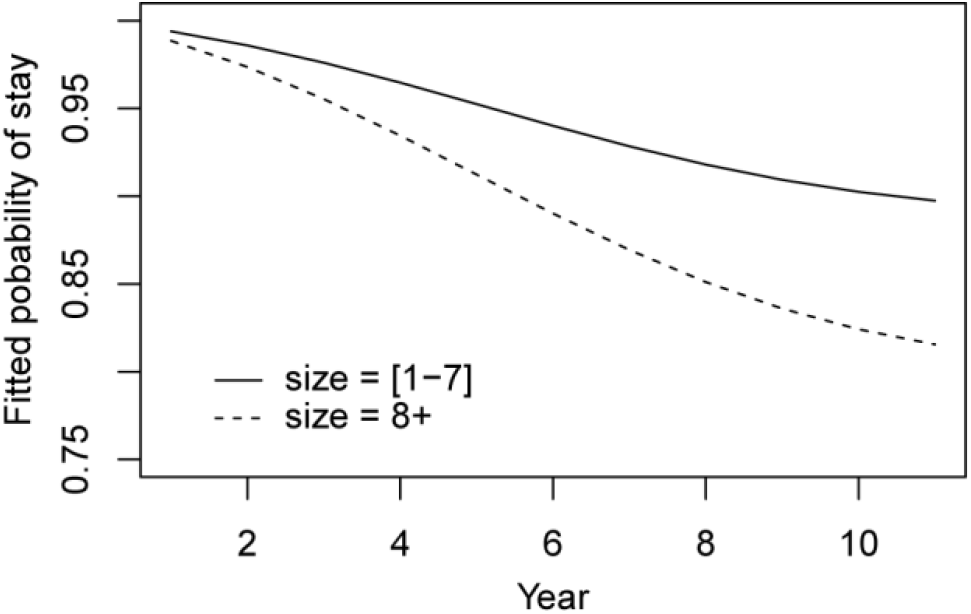
Household size and predicted probability of staying at one’s duolocal household. The predictors are individuals’ sex as time-invariant variables, and age, average relatedness to others in the households, household size, year, and year square as time-varying variables. In x-axis, years 1 – 11 = the year 2007 to 2017. Individuals living in households larger than 7 had a higher hazard of dispersal (solid line) than those in smaller households (dashed line).

### 3.4. Reproduction and mortality in matrilineal Mosuo populations

In the year 2007, there were on average 2.13 reproducing-age females (15-50 years old, range 0–7, s.d. = 1.22) and 2.38 reproducing-age males (15-50 years old, range 0–8, s.d. = 1.48) in each duolocal family (n = 378), and these numbers hardly changed in following 10 years (Fig. S5). In families smaller than 8, when family-group size increased by one, the annual probability of birth of reproducing-age females in these families decreased by 23.5% (HR = 0.745; 95% CI = [0.573, 0.979], n = 2,610 person-years, 91 events; Fig. 5 and Table S4); whilst in families larger than 6, family-group size had no significant influence on annual probability of birth (n = 4,951 person-years, 183 events, Table S4). These results show some support for prediction 2, in the sense that females in small families were more likely to reproduce than those in big families. Moreover, they imply that most Mosuo duolocal families were not at optimum, and the optimal group size should be smaller than the mean, which might be the stable group size (Shen et al., 2017; Sibly, 1983). Besides, although permitted by government policy, females in duolocal households were reluctant to give birth to a third child. The hazard of giving birth to females who already had two alive children was only 2.8% or 11.7% of that of females who had no child (Table S5).

**Fig. 5.**
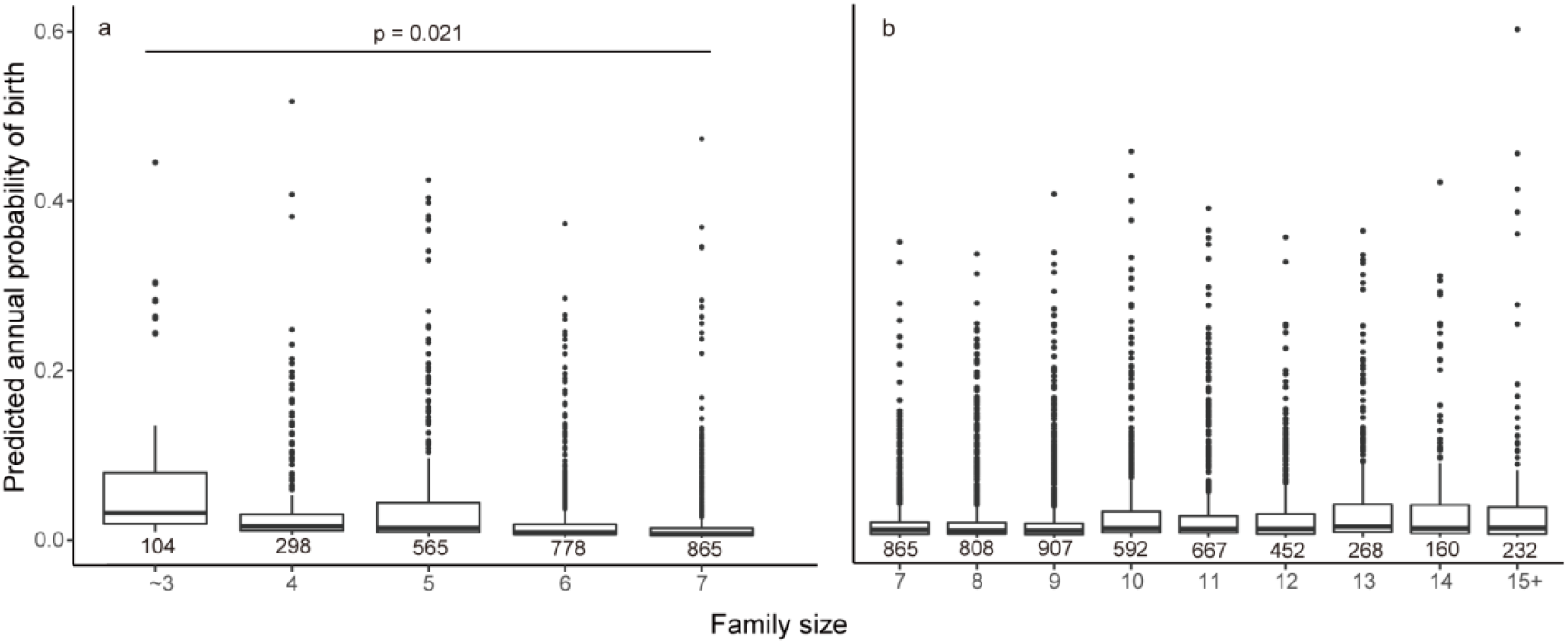
Boxplot of model-predicted values for the annual probability of birth obtained from a mixed complementary log-log link model for different family sizes. The predictors are the female’s ID as random variables, and household size, age (centered), and alive offspring number as time-varying variables (fixed effects). P values are shown where there were significant differences between family sizes, and numbers below each box were the number of person-years. (a) In families ≤ 7, the annual probability of birth was negatively correlated to family size. (b) In families ≥ 7, the annual probability of birth of reproducing-age females was not associated with family size.

We predict that the probability of females giving birth to a child decrease as family size arises, whist mortality increases; in households with a size smaller or larger than the threshold, females have fewer offspring, longer birth interval, latter onset, and earlier termination of birth. We measured how the size of family groups related to birth intervals, age of first birth, number of alive offspring, and co-residing offspring of reproducing-age females, in 2007. The results show some support for our prediction (Table S5). For families ≤ 7, females in larger families started birth earlier (ES = -0.522, 95% CI = [-0.897, -0.147]), whilst for families ≥ 7, females in larger families started birth later (ES = 0.200, 95% CI = [0.057, 0.343]). For families ≤ 7, the number of alive offspring and co-residing offspring of females living increased as group size increased (ES = 0.108 and 0.140, 95% CI = [0.009, 0.21] and [0.057, 0.222] for the number of alive and co-residing offspring, respectively), and for families ≥ 7, they non-significantly but negatively related to group size (ES = -0.018 and -0.019, 95% CI = [-0.047, 0.011] and [-0.05, 0.011] for the two, respectively). Birth intervals were not related to group size, but the estimates were also in a favourable direction (ES = -0.126 and 0.014, 95% CI = [-0.392, 0.140] and [-0.077, 0.105] for families ≤ 7 and ≥ 7, respectively).

As current family size might reflect the past family size, we also tested how five aspects of reproduction of elder females aged 40+ from these families changed with household size in year 2007, but we found no supportive evidence for current family size affecting reproduction of elder females (see SI text1 and Table S6). Similarly, we found no evidence for the death of family members influenced by family-group size (discrete-time event history analysis, p = 0.427 and 0.921 for individuals in families ≤ 7 and ≥ 7, respectively). Older members (HR for age = 1.09 and 1.084, 95% CI = [1.074, 1.108] and [1.073, 1.096] for families ≤ 7 and ≥ 7, respectively) or men (HR for sex (women as reference) = 0.378 and 0.519, 95% CI = [0.231, 0.608] and [0.368, 0.725] for families ≤ 7 and ≥ 7, respectively) were more likely to die. However, it should be noted that mortality in this location is low.

### 3.5. Within-household cooperation in matrilineal Mosuo populations

On average, Mosuo men from duolocal households contributed more money in the economic game to their households than women. Men contributed 8.299 ± 1.888 to their households, and women contributed 7.205 ± 2.423. Results from a multiple-level model showed that there were significant sex differences (Fig. 6). Although household size or its interaction with average relatedness to others did not significantly affect an individual’s contributions (ES for size = 0.025; 95% CI = [-0.071, 0.123]), the interaction was in a favourable direction (ES = -0.777; 95% CI = [-1.607, 0.052]), where an increase of relatedness raised less contribution for individuals in larger households, relative to those in smaller ones. Besides, age positively interacted with average relatedness with other family members (ES = 0.183; 95% CI = [0.001, 0.372]). That is, higher relatedness led to increased contribution for the elders more than for the younger ones.

**Fig. 6.**
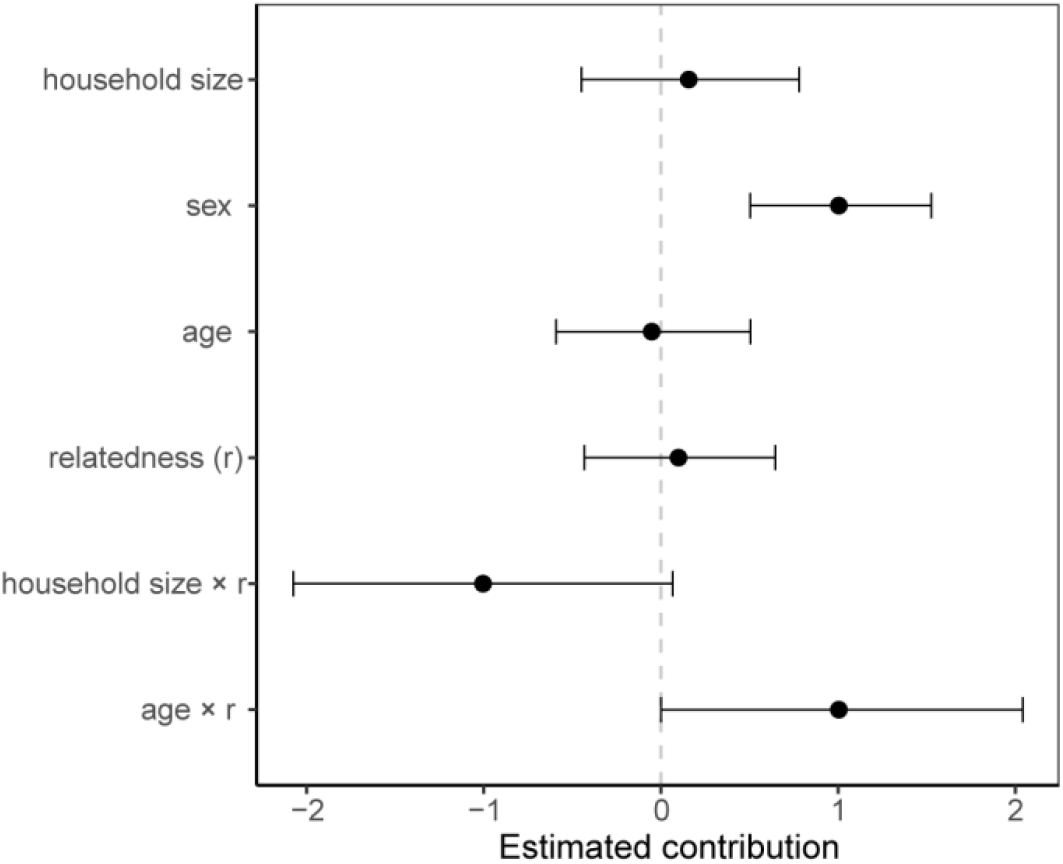
**Estimates predicting contribution to the household. Estimates were** standardized over 2 s.d. to allow comparison between continuous and binary predictors (Gelman, 2008). Error bars are 95% confidence intervals. Intercept (ES = 7.220; 95% CI = [6.833, 7.629]) not shown for clarity.

We also looked at how group size influenced the proportion of family members who worked on the farm (Fig. S6), and we found a small but significantly negative effect of large size on cooperation level for families smaller than 8 (size = 1-7, with size ≤ 3 combined). In families smaller than 8, when the group size increased by one, the proportion of working members decreased by 3.33% (ES = -0.033, 95% CI = [-0.064, -0.002]). Whist in families larger than 6 (size = 7-19, with size ≥ 15 combined), group size did not relate to the proportion of working members on the farm.

## 4. Discussion

Group size is an important structural characteristic of social groups, including human families, which might influence the relatedness and interactions of members (Rodrigues & Gardner, 2013b). Household size has decreased globally (Powell et al., 2018; United Nations Department of Economic and Social Affairs Population Division, 2017, 2019). Similar patterns were observed in our study area. Lower birth rates, higher rates of dispersal, and market economy, which lead to a decrease of duolocal residence (Ji et al., 2016; Mattison, 2010) and matrilineal households in this area (Walsh, 2004), might account for the observed decrease in family size. In this paper, we investigated the effects of family size on the average inclusive fitness of households, using evolutionary game models among co-residing relatives. Furthermore, we measured how the size of Mosuo duolocal families changes in a rural Chinese population in a decade, and how the change affects the average relatedness of these families, female reproduction, dispersal, and helping behavior of the co-residing family members.

Average relatedness (or local relatedness) has been proven to be important in determining cooperation and competition pattern among group members (Bowles & Posel, 2005; Brent et al., 2015; Cant & Johnstone, 2008; Croft et al., 2021; Johnstone & Cant, 2010; J. Wu et al., 2013). However, the relationship between group size and local relatedness has been less well explored (Croft et al., 2021). Our empirical data showed that the average relatedness of family-group members decreased with the increase of size in a duolocal household, while in other types of post-marital residences, such as neolocal, patrilocal, or matrilocal households, the averaged relatedness of these families increased and then decreased with the increase of family size. It should be noted that in theory, the increasing household size of a nuclear family (or neolocal household, which only contains husband, wife, and their children) would always lead to the average relatedness rising towards 0.5. Nevertheless, our empirical data from neolocal households showed a decreased average relatedness when a family is getting bigger than 4 (Fig. 2b), implying that a nuclear family might rarely grow too big unless the family accepts some immigrants under certain circumstances. In consequence, with the same costs and benefits of interaction between co-residence members, individuals living in a large family might infer a less inclusive fitness advantage than those in a smaller family. Our model predicted that there should be an evolutionarily-stable family size, given any combinations of costs and benefits. This conclusion applies to family groups of all kinds of kinship systems and post-marital residence, as long as the negative correlation between family size and average relatedness holds.

The empirical data from duolocal Mosuo provided evidence to support our model predictions. The onset of birth of Mosuo females was earliest, and the alive offspring number was highest for those living in mean-size family groups, relative to those in groups of larger or smaller sizes. Annual probability of birth for reproducing-aged women negatively related to family size in households with a size above average, whilst no relationship was found for those in larger households. These results imply that Mosuo women living in small family groups are more likely to give birth to a new baby (reached the peak at size ≤ 3 rather than mean group size). We found no relationship between current family size and reproduction of elder women or mortality of family members, however, the real factor that affects the reproduction of elder females should be group size in their reproducing age (which we do not know), rather than current group size. We found that larger duolocal households were less likely to tolerate immigration. Mosuo in larger family groups (> = 7) experienced higher hazards of dispersal than those in smaller ones, and people who are less related to other members of households were more likely to disperse. These results are consistent with our previous study, which found that a large number of adult siblings drove Mosuo men and women out of their natal household (He et al., 2016) Our study also found few dispersals of elder men or women during a decade. These results suggest that the younger generation either has lower relatedness with other members of the household, or has more opportunities than the elders to establish a new household in this area or the cities, or both.

Whilst our previous work found that married Mosuo men work less on the farm, we found Mosuo men in duolocal households contribute more to their households than women, in line with our other game results where men showed a higher level of cooperation with villagers (J.-J. Wu et al., 2015). Our result might reflect a tendency for Mosuo men rather than women to signal their cooperativeness, or Mosuo men tending to help their household in the form of monetary contribution (rather than working on a farm or doing housework) in everyday life.

Rodrigues and Gardner (Rodrigues & Gardner, 2013b) predict that temporally group size decreasing can favour harming others in the group. Although group size decreased slightly in the last ten years in our study area, we still found a considerable high level of cooperation within households. Mosuo men and women contribute more than 8 and 7 out of 10 to their household respectively, whilst their average donation to villagers in a dictator game was lower than 5 (J.-J. Wu et al., 2015). Although we didn’t find evidence that group size directly influenced the contribution to a duolocal household by Mosuo men and women in an economic game within families, the negative effect of interactions between group size and relatedness which had marginal significance, suggested that household size decreased the positive effects of relatedness on cooperation between family members to some extent, in consistent to the hypothesis that the limited dispersal that increase the relatedness between individuals also lead to intensified competition among kin (Hamilton, 1964a, 1964b; Queller, 1992; West et al., 2002). The proportion of members working on the farm was negatively related to group size in households smaller than eight. Moreover, higher relatedness to other group members had larger effects on elders’ contribution than youngers’, which might suggest a higher conflict level in the younger generation than elder ones. These results highlight the importance of elder generations in maintaining social cohesion in large families, as noted in other anthropological contexts (Chagnon, 1979), and that kin selection affected within-household cooperation (Hamilton, 1964a, 1964b).

The family-size dynamic is not only influenced by but also affects the demographic events, and it also influences the interactions and ultimately inclusive fitness of the family members (through changing of local relatedness (Croft et al., 2021), possibly also different costs and benefits of helping co-residing members). Although there are many advantages to living in large families (Chen et al., 2023; Chomtohsuwan & Nagashima, 2010b; Dessy et al., 2021b; Sauerborn et al., 1996; Thomas et al., 2018), there may also be drawbacks, such as decreased cooperation and an increase in conflict due to decreasing average relatedness. For example, it has been reported that more effort should be spent on reproductive competition between residing father and son and between brothers under polygyny than monogamy (Ji et al., 2014), and Tanzanian children from polygynous families (bigger households) show poorer growth performance than those from monogamous families (smaller households) (Hadley, 2005). Our theoretical model predicted the existence of an evolutionarily stable household size that households should maintain given any costs and benefits. Longitudinal data analyses support that average relatedness is associated with helping behavior in the Mosuo duolocal household, and these households tend to keep a higher level of average relatedness, with individuals being more likely to emigrate from and less likely to immigrate into large families. These results should apply to families of other types of post-marital residence, especially extended families where the negative relationship of average relatedness and group size most likely exists. These findings are helpful to understand the variation in family size in different cultures and demonstrate the importance of kin selection in regulating family dynamics.

## Funding

This work was supported by the National Natural Science Foundation of China (grant numbers 31971401, 31971403, 32071610, and 31971511) and the European Research Council (grant number EvoBias AdG 834597).

## Acknowledgements

We thank members of TEG in IOZ and Professor Hua Wu, Dr. Zhen-Hua Luo, and their students from Central China Normal University, Zhizhi A’ge, Ya-Juan Liu, Youzhuacier Wabu, Erchema A’wa, Bima Yang, Kumu A’ge, Ruozuocier Wabu, Caiwenglongbu Li, and HEEG in Lanzhou university for helping in collecting data. We thank Mosuo for their participation in this study.

## Declaration of interests

The authors declare no interest conflict.

## Notes

### Competing Interest Statement

The authors have declared no competing interest.

