## Supplementary Information for "Kin selection underpins family dynamics in rural China"

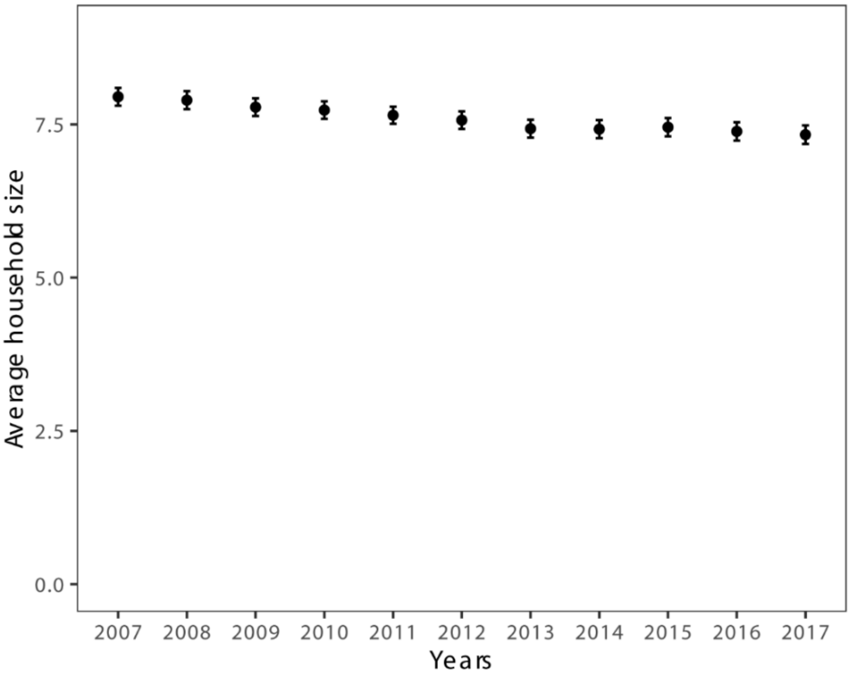


**Fig. S1. Average household size slightly decreased during 2007 and 2017**. Mean size of 378 households, labeled as duolocal in year 2007, decreased among year 2007 (mean size 7.95 ± 2.84) and 2017 (mean size 7.33 ± 2.96). Mixed effect model found significant negative relationship of group size and years (ES = -0.063, 95% CI = [-0.073, -0.053], household id as random effect).

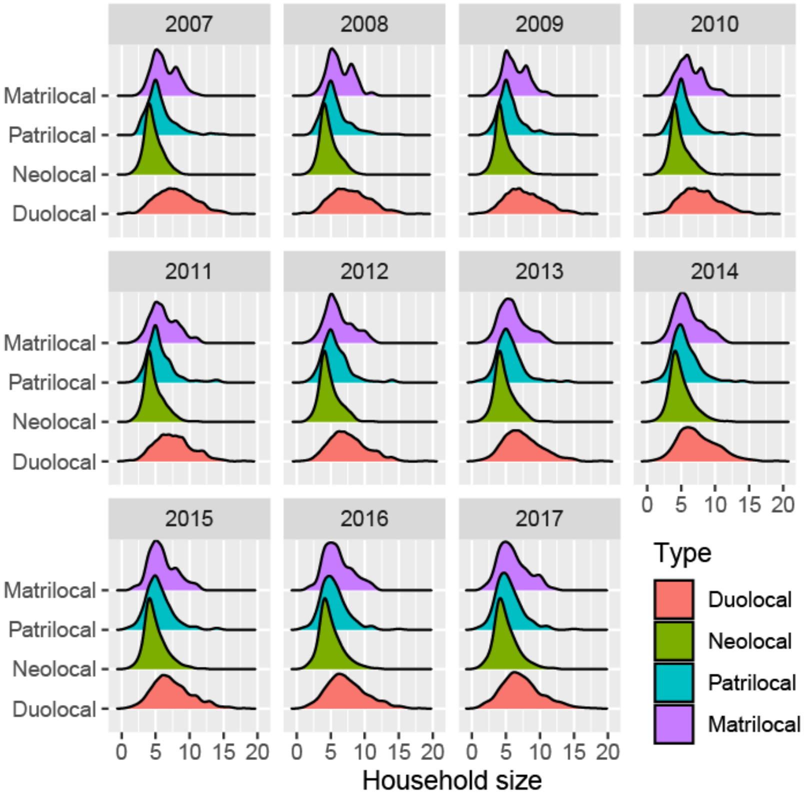


**Fig. S2. Distribution of family-group size in a decade**.

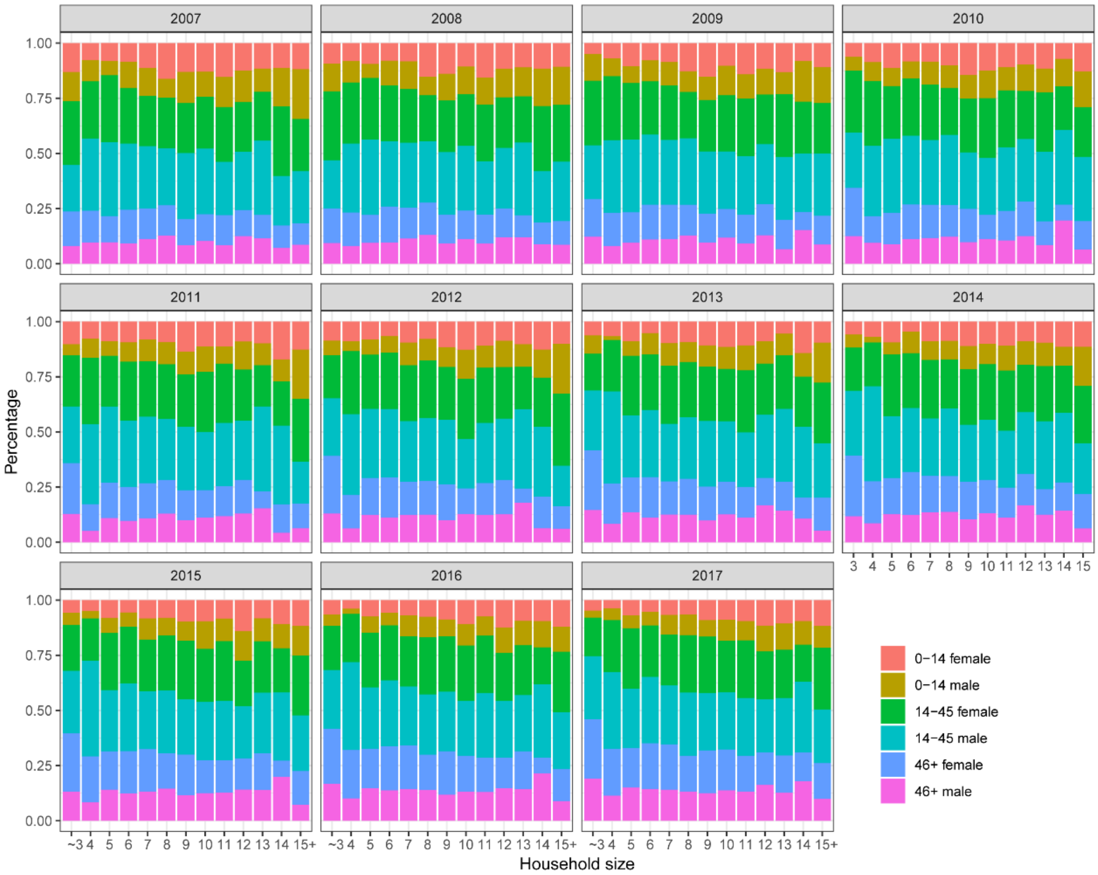


**Fig. S3. Composition of household did not vary with group size during 2007 and 2017.**

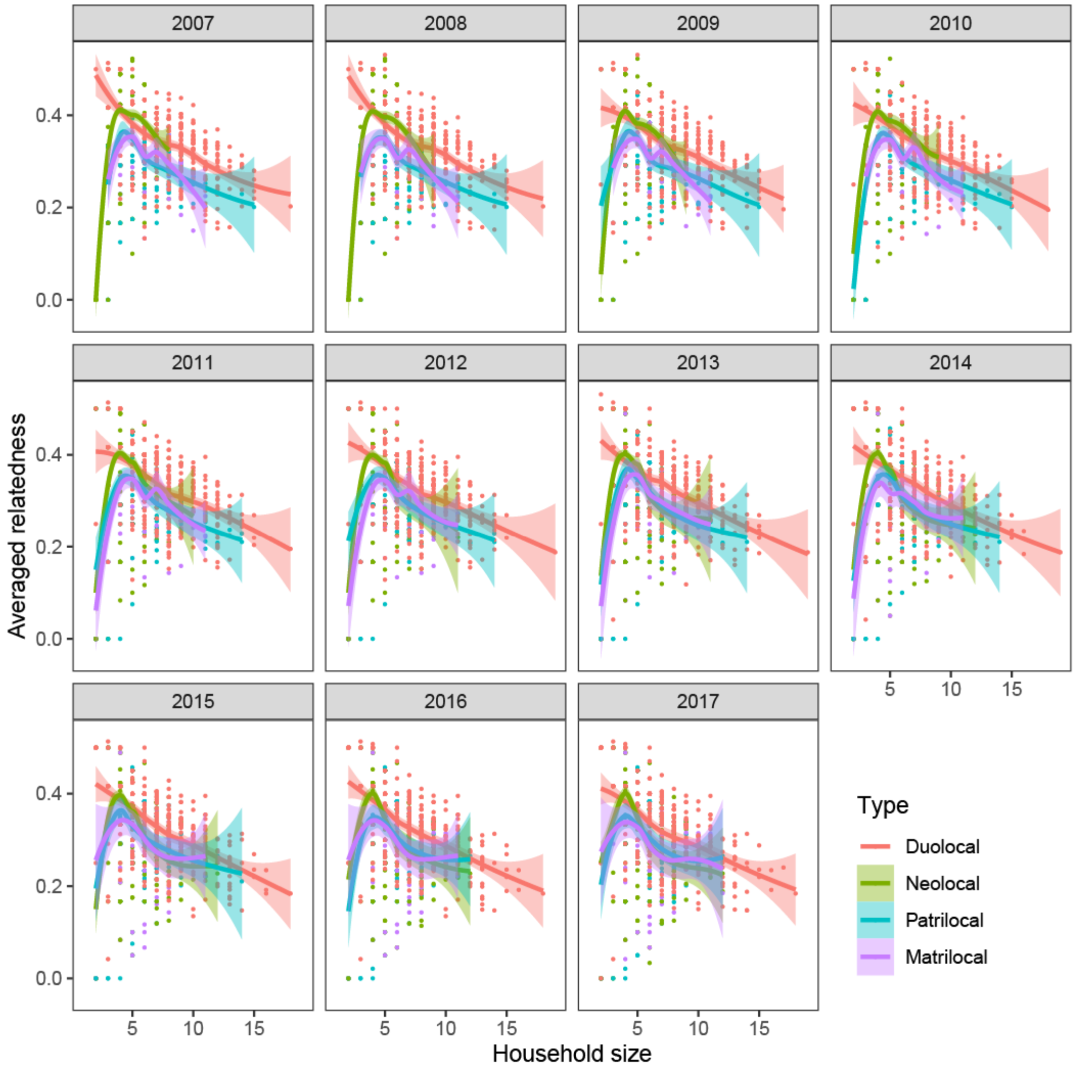


**Fig. S4. The relationship between averaged relatedness and household size of households, from year 2007 to 2017.** Average relatedness of duolocal households decreased with group size, and it increases first and declines latter as the sizes of other three types of household increase. This pattern holds for all the 11 years. Note that we defined the type of households according to the residence type applied by most married members in 2007. Curves fitted using loess. Shaded areas indicate 95% confidence interval.

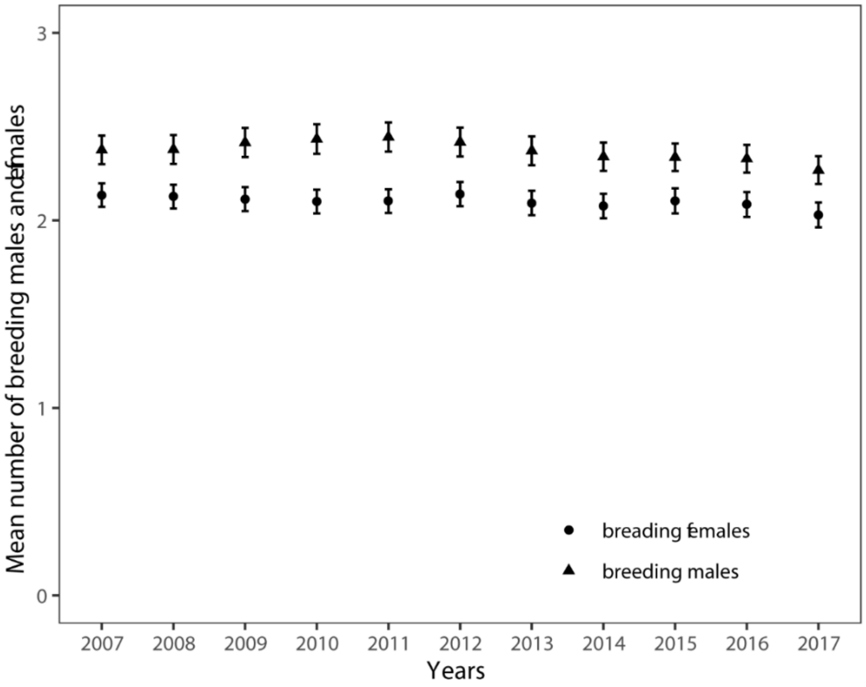


**Fig. S5. Changes of mean numbers of reproducing-aged females and males in 378 Mosuo households**. In year 2017, each household contained a mean of 2.03 (age 15-50, range 0-7, s.d. = 1.29) reproducing-age females and 2.27 (age 15-50, range 0-8, s.d. = 1.44) reproducing-age males, whilst in year 2007 there were 2.13 (age 15-50, range 0-7, s.d. = 1.22) females and 2.38 (age 15-50, range 0-8, s.d. = 1.48) males respectively. Error bars indicate mean ± s. e. Circle points, reproducing-aged females; Triangle points, reproducing aged males.

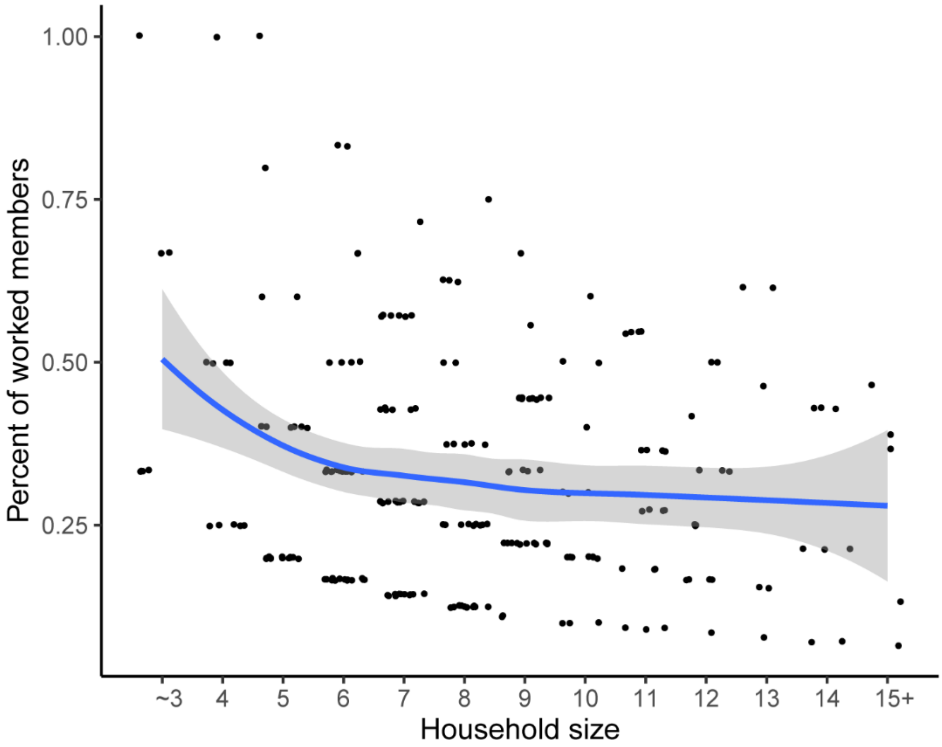


**Fig. S6. Observed proportion of members worked in farm and group size**.

**Table S1. The changes of average household relatedness after each demographic event.** Δr stands for changes of relatedness; s.d. indicates standard deviance. Different letters ([a - d]) indicate significant differences (Dwass-Steele-Critchlow-Fligner test, $p<0.01$).

| Events | n | Mean Δr | s.d. |
|---|---|---|---|
| Death | 200 | 0.016 [a] | 0.035 |
| Emigration | 463 | 0.010 [b] | 0.040 |
| Birth | 303 | -0.025 [c] | 0.034 |
| Immigration | 142 | -0.054 [d] | 0.073 |
| Total | 1108 | -0.013 | 0.046 |

**Table S2. The effect of household size on annual probability of dispersal for adults in duolocal households (n = 15,967 person-years; events = 275)**. Intercept (HR = 0.006, 95% CI = [0.003, 0.012]) and time variable (year (1.458, [1.224, 1.748]) and year square (HR = 0.968, 95% CI = [0.953, 0.983])) not shown for clarity. Statistical significance indicates in bold; HR stands for hazard ratio, and CI for confidence intervals.

| Variables | HR [95% CI] | P value |
|---|---|---|
| Fixed effects | | |
| **Family size (ref: 1-7)** | | |
| **≥ 8** | 1.884 [1.357, 2.668] | **<0.001** |
| **Age (scaled)** | 0.804 [0.706, 0.912] | **0.001** |
| **Relatedness to others** | 0.189 [0.047, 0.793] | **0.021** |
| **Sex (ref: men)** | | |
| **Women** | 1.357 [1.052, 1.753] | **0.019** |
| Random effects | Variance (s.d.) | |
| Household ID | 0.753 (0.868) | |
| n | 378 households | |

**Table S3. Number of immigrations happened in a specific year and group size (n = 4,153 household-years, 378 households; events = 143 happened in 106 household-years) among year 2007 and 2017**. Intercepts (ES = 0.06, 95% CI = [0.038, 0.083]) not shown for clarity. Family size ≤ 3 and ≥ 15 were combined. CI stands for confidence intervals.

| Variables | Estimate | 95% CI |
|---|---|---|
| Fixed effects | | |
| **Family size** | **-0.003** | **-0.006, -0.001** |
| Random effects | Variance (s.d.) | |
| Household ID | 0 (0.018) | |
| n | 378 households | |

**Table S4. Annual probability of birth from mixed complementary log-log link models.** Intercepts (HR = 0.101, 95% CI = [0.015, 0.422] for model 1; HR = 0.017, 95% CI = [0.006, 0.04] for model 2) not shown for clarity. Family size ≤ 3 and ≥ 15 were combined. Statistical significance indicates in bold; HR stands for hazard ratio; CI stands for bootstrap confidence intervals.

| Variables | Model 1: family size < = 7 | | Model 2: family size > = 7 | |
|---|---|---|---|---|
| | HR [95% CI] | P value | HR [95% CI] | P value |
| Fixed effects | | | | |
| **Family size** | **0.745 [0.573, 0.979]** | **0.021** | 1.065 [0.972, 1.146] | 0.172 |
| **Age (centered)** | 1.058 [1.003, 1.112] | 0.075 | **1.094 [1.064, 1.128]** | **<0.001** |
| Alive offspring number (ref: none) | | | | |
| One | 0.772 [0.325, 1.753] | 0.566 | 1.403 [0.898, 2.363] | 0.255 |
| **Two** | **0.117 [0.031, 0.353]** | **0.002** | **0.028 [0.011, 0.069]** | **<0.001** |
| Random effects | Variance (s.d.) | | | |
| Female ID | 3.16 (1.9) | | 2.218 (1.489) | |
| n (females) | 338 | | 595 | |

**Table S5. The effect of household size on reproduction of reproducing-age females (age = 15 - 50) of duolocal households in year 2007**. A series of linear regressions were done for birth interval, first birth age, and co-residing offspring number, and two Poisson regressions for alive offspring number, all the models with household size as independent variable. Four models for offspring number also controlled for age square (standardized age and its square), and others controlled for females' age (centered by mean). Family size ≤ 3 and ≥ 15 were combined. Statistical significance indicates in bold. CI stands for confidence intervals.

| Dependent variables | Family size < = 7 | | Family size > = 7 | |
|---|---|---|---|---|
| | Estimate (Sig.) [95% CI] | n | Estimate (Sig.) [95% CI] | n |
| Birth interval | -0.126 (0.35) [-0.392, 0.14] | 173 | 0.014 (0.759) [-0.077, 0.105] | 437 |
| **First birth age** | **-0.522 (0.007) [-0.897, -0.147]** | **205** | **0.2 (0.006) [0.057, 0.343]** | **504** |
| **Alive offspring number** | **0.108 (0.036) [0.009, 0.21]** | **272** | -0.018 (0.227) [-0.047, 0.011] | 624 |
| **Co-residing offspring** | **0.14 (0.001) [0.057, 0.222]** | **272** | -0.019 (0.204) [-0.05, 0.011] | 624 |

SI Text 1

We also measured how family size varied with number of adult offspring and co-residing offspring, age of first birth, age of last birth, and average birth interval of females over 40 years old in these households, who had completed most reproduction. We modeled the relationship between these variables and family size in a series of linear regressions and Poisson regressions, with age of females controlled, and sample sizes varied with specific regressions (Table S5). We combined family size ≤ 3 and ≥ 15, for families with size smaller than 3 or larger than 15 were rare. Analysis of birth interval only involved females had more than one child. We found no significant effect of current family size on their age of first birth, age of last birth, or number of adult offspring living to age 15 and over. Group size positively associated with number of co-residing offspring (ES = 0.201, 95% CI = [0.072, 0.329] for families ≤ 7, and ES = 0.07, 95% CI = [0.009, 0.132]), although the latter is more likely a cause rather than a result of size increasing. Contrary to our prediction, in families larger than 6, birth intervals of elder females negatively related to household size (ES = -0.11, 95% CI = [-0.198, -0.021]), though no significant relationship was found in a model of those smaller families.

**Table S6. The effect of household size on reproduction of females over 40 years old of duolocal households in year 2007.** A series of linear regressions were done for birth interval, first and last birth age, and co-residing offspring number, and two Poisson regressions for adult offspring number (living to age 15), all the models with current household size as independent variable. Four models for offspring number also controlled for age square (standardized age and its square), and others controlled for females' age (centered by mean). Family size ≤ 3 and ≥ 15 were combined. Statistical significance indicates in bold. CI stands for confidence intervals.

| Dependent variables | Family size < = 7 | | Family size > = 7 | |
|---|---|---|---|---|
| | Estimate (Sig.) [95% CI] | n | Estimate (Sig.) [95% CI] | n |
| **Birth interval** | 0.183 (0.215) [-0.107, 0.473] | 155 | **-0.11 (0.016) [-0.198, -0.021]** | 329 |
| First birth age | 0.067 (0.749) [-0.348, 0.483] | 167 | 0.09 (0.337) [-0.094, 0.273] | 359 |
| Last birth age | 0.323 (0.326) [-0.324, 0.969] | 167 | -0.009 (0.946) [-0.260, 0.242] | 359 |
| Offspring living to age 15 | 0.028 (0.468) [-0.046, 0.104] | 175 | 0.017 (0.197) [-0.009, 0.042] | 386 |
| **Co-residing offspring** | **0.201 (0.002) [0.072, 0.329]** | **175** | **0.07 (0.026) [0.009, 0.132]** | 386 |
